# Cervical Spinal Cord Modulation with Repeated Epidural Stimulation in Healthy Adult Rats

**DOI:** 10.1101/2022.07.11.498751

**Authors:** Pawan Sharma, Hema Rampersaud, Prithvi K. Shah

**Affiliations:** Division of Rehabilitation Sciences, Department of Physical Therapy, School of Health Technology and Management, Stony Brook University, Stony Brook, New York 11727; Department of Physiology and Biophysics, Stony Brook University, Stony Brook, New York 11727

## Abstract

The effects of spinal epidural stimulation (ES) in regaining various physiological functions after a spinal cord injury (SCI) are well documented. Spinal evoked motor responses (SEMR) are commonly utilized experimental outcomes in longitudinal pre-clinical and human studies to reflect the in-vivo physiological changes in neural networks secondary to a neurological insult or neuro-rehabilitative treatments utilizing ES. However, it remains unknown if the repeated exposure to ES during SEMRs testing itself modulates the spinal cord physiology and hence the SEMRs characteristics. To address this issue, cervical ES was delivered to the healthy adult rat’s cervical cord using standard stimulation paradigms during multiple sessions (∼17 hours that spanned across 100 days). Cervical SEMR and electromyography (EMG) activity from forelimb muscles during a reaching & grasping task were collected before and after 100 days. We noted persistent increase in the cervical SEMR and forelimb muscle activity during reaching & grasping task relative to baseline at the end of the stimulation period indicating increased spinal and cortical excitability. Findings from the present work suggests that cervical SEMRs are amenable to modulation by routine ES testing protocols, with prominent changes in the mono and poly synaptic component of evoked responses. Additionally, since multiple testing sessions of cervical ES alone increases the excitability of the intact spinal cord, we suggest that SEMR data be used with caution to infer the physiological status of the spinal circuitry in longitudinal studies involving multiple SEMR testing sessions. Our findings also recommend involving appropriate control groups, motor behavior correlates, and practicing caution while utilizing normalization methods to allow meaningful functional interpretation of SEMR profiles following a SCI.

## Introduction

Epidural stimulation (ES) has shown to be effective in improving several physiological functions, such as independent standing, stepping, and autonomic functions, following spinal cord injuries (SCI) (Angeli *et al*., 2018; Harkema *et al*., 2018; Darrow *et al*., 2019). Pre-clinical studies investigating the effectiveness of ES in recovering sensorimotor function post-SCI often employ spinal evoked motor responses (SEMR) as a biomarker of recovery (Gad *et al*., 2015; Alam *et al*., 2017). SEMRs, recorded as electromyography (EMG) signals from muscles secondary to spinal ES (Gerasimenko *et al*., 2006; Sharma & Shah, 2021b), are physiological readouts of neural activity in the intact or injured spinal cord and are suggested to correlate with motor recovery following a SCI (Lavrov *et al*., 2008; Hofstoetter *et al*., 2019; Militskova *et al*., 2020). Longitudinal studies investigating ES effectiveness in sensorimotor recovery following a SCI in rats involve multiple SEMR data collection sessions (2-4 times in a given experiment) (Gad *et al*., 2015; Alam *et al*., 2017), with each session ranging between 45-60 minutes and involves delivering ES of various intensities and frequency (based on previous experience and ongoing work in the lab).

Recent work involving lumbar ES for 15 minutes in healthy adult rats demonstrated an increase in the amplitudes of the obtained SEMR, with effects lasting for eight minutes post-stimulation (Taccola *et al*., 2020). Similar facilitatory effects have been demonstrated in healthy humans with 10-30 minutes of cervical and lumbar transcutaneous stimulation. Immediate gains in the obtained spinal and cortical evoked responses and improved motor output suggest increased modulation susceptibility of the spinal circuitry even with short bouts of electrical stimulation (Benavides *et al*., 2020; Kumru *et al*., 2021). These observations raise an important question: will discrete and independent bouts of ES that are not aimed for therapeutic effects but employed in stimulation testing protocols also modulate SEMRs? Addressing this question has implications for interpreting SEMR data in longitudinal studies which employ repeated stimulation testing protocols. Addressing whether modulation of SEMRs, if any, is a consequence of repeated stimulation during SEMR data acquisition per se or an outcome of an independent variable in a study will allow practicing precautions while collecting, analyzing, and interpreting SEMR data in longitudinal study designs involving regular SEMR data acquisition.

To address the posed research question, data collected as a part of the larger project aimed at identifying the potential neural components interacting with cervical ES in healthy adult rats were explored (Sharma & Shah, 2021a). Rats underwent a series of stimulation paradigms involving ES delivery targeting C6 and C8 cervical segments using stimulation protocols commonly employed in longitudinal pre-clinical studies (Courtine *et al*., 2009; Alam *et al*., 2017) for approximately seventeen hours and spanning over 100 days. The findings from the present work suggest that cervical SEMRs are amenable to modulation by routine ES testing protocols, with prominent changes in the mono and poly synaptic components of evoked responses. Additionally, repeated stimulation results in increased muscle activation of select forelimb muscles during the reaching & grasping motor task.

## Methods

### Ethical Approval

The study was carried out in accordance with the recommendations of the National Institutes of Health Guide for the Care and Use of Laboratory Animals. The protocol was approved by the Stony Brook Office of the Vice President for Research, and Institutional Animal Care and Use Committee (IACUC). Rats were obtained from Charles River Laboratories (Charles River Laboratories International Inc., U.S.A) and were housed in standard conditions (26 ± 1 °C; 12 h−12 h dark-light cycle) with access to standard food and water ad libitum.

### Experimental Design

An outline of the experimental design is shown in **figure 1-a**. A total of seven adult female Sprague-Dawley rats (240-260 grams body weight) were trained for reaching & grasping performance with the preferred paw after initial handling and acclimatization. Following training, rats underwent surgical procedures for implantation of chronic electrodes into the muscle (for electromyography, EMG) and the spinal cord (for epidural stimulation, ES). Eleven days following recovery from implant surgeries, rats received sessions of ES during cervical SEMR data acquisition spanning over 100 days while awake.

**Figure 1:**
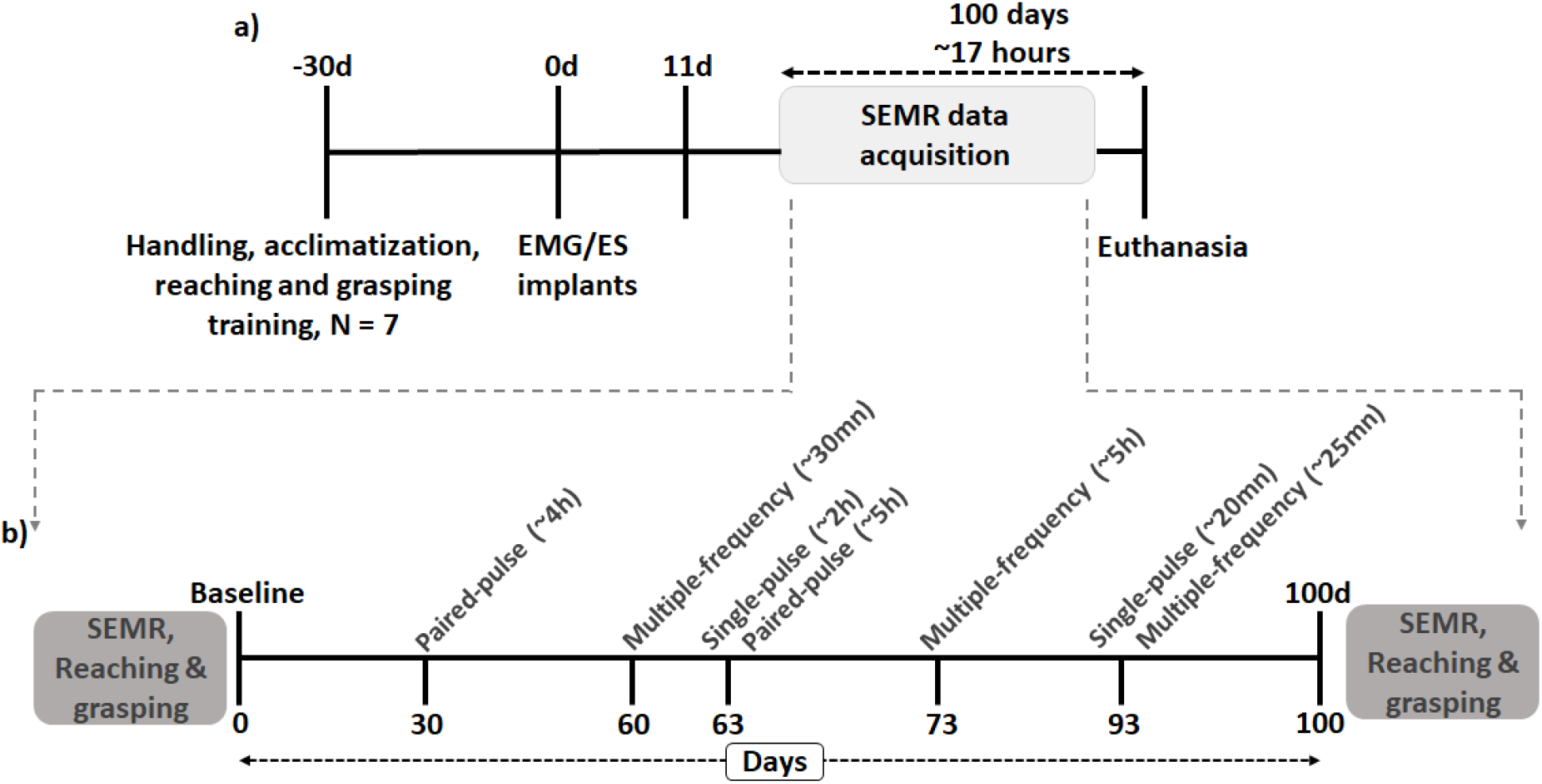
Experimental design and stimulation sessions: **a)** All rats were handled, acclimatized and trained in a reaching & grasping task before undergoing surgical procedures. Rats then underwent chronic electromyographic (EMG) implantation surgery in selected forelimb muscles and cervical epidural stimulation (ES) electrode implantation at C6 and C8 segments. Following recovery from surgeries at around eleven days, rats received cervical ES of different stimulations protocol during spinal evoked motor responses (SEMR) data acquisition sessions spread across multiple sessions over 100 days for a total of ∼17 hours. At the completion of these experiments, all rats were euthanized using transcardial perfusion. **b)** During stimulation sessions, rats received ES of varying stimulation protocol (singe-pulse, paired-pulse, multiple frequency) while awake. Numerals in parenthesis represent the average duration of stimulation in hours (hr) or minutes (mn) per rat for the specific stimulation protocol during each stimulation session. Cervical SEMR secondary to single-pulse bipolar ES and reaching & grasping task were collected before (baseline) and end of the whole stimulation period (100d).

### Reaching & grasping training

Reaching & grasping training was performed as described previously (Whishaw & Pellis, 1990; Whishaw *et al*., 2010; Sharma & Shah, 2021b). Briefly, following dominant paw identification, each rat underwent Reaching & grasping training until they reached a seventy percent success rate during the task. During each training session, a total of thirty sugar pellets (45 mg banana-flavored sugar pellets 0.14” diameter, Dustless Precision Pellets, Bio-Serv, Frenchtown, NJ, USA) were provided, and successful Reaching & grasping attempts were recorded to calculate the success rate. Throughout the habituation and training period, all rats were put on a food restriction protocol wherein they received a standard laboratory diet equivalent to 5% of their body weight daily to maintain approximately 90% of their initial weight (Whishaw & Pellis, 1990; Whishaw *et al*., 2010; Zemmar *et al*., 2015).

### EMG and ES electrode implantation surgical procedure

All surgeries were performed under aseptic conditions with rats deeply anesthetized with isoflurane gas (1.0-2.5% via facemask as needed) on a heated table to maintain body temperature at 37^0^C as described previously (Rascoe *et al*., 2018; Richards *et al*., 2019). After surgery, rats were placed in an incubator until fully recovered and administered antibiotics (Enrofloxacin, 40 mg/kg) and analgesics (Buprenex, 0.03 mg/kg) as needed for up to 5 days. Thereafter, rats were housed singly in cages to avoid damaging each other’s head plugs through social interaction.

### EMG and ES electrode implantation

For EMG electrode implantation, select forelimb muscles (flexor digitorum superficialis (FDS), extensor digitorum (ED), pronator (PR), deltoid (DL)) of the preferred paw were implanted with intramuscular recording electrodes. Briefly, following skull incisions, one connector (Part # A22004-001, Omnetics Connector; MN, USA) with Teflon-coated stainless-steel wires (Part # AS632 and AS631; Cooner Wire Co, Chatsworth, CA, USA) was attached securely to the skull with screws and dental cement. Skin and fascial incisions were made to expose the belly of the selected muscles. Bipolar intramuscular EMG electrodes were formed by stripping ∼1mm Teflon coating from the stainless-steel wires and secured into the mid-belly of each muscle. After electrode implantation into the muscle, proper placement of EMG electrodes into the desired muscle was verified by stimulating the muscle via the head plug connector.

For ES electrode implantation, following longitudinal skin incision between C2 and T2 spinous process and retraction of underlying fascia and upper back muscles, a partial at laminectomy C4 and a complete laminectomy at C5 and C7 was performed. Laminectomies at C5 and C7 exposed the spinal cord segments C6 and C8, respectively. A small region (∼1 mm notch) of the Teflon coating was removed from each wire to form the stimulating electrodes that were then secured to the dura at the midline of the spinal cord at each site with 8.0 Ethilon sutures (Rascoe *et al*., 2018; Sharma & Shah, 2021a).

Approximately one inch of Teflon coating was stripped at the distal end of an additional wires and placed subcutaneously near the inferior scapular angle to serve as a reference electrode for EMG and ES electrodes. Post-mortem, position of all electrodes were verified to ensure the integrity and proper placement of electrodes.

### Stimulation and recording procedures

As the main objective of the present was to investigate the neuromodulatory effects of repetitive ES during SEMR data collection on the intact spinal cord, we used varied stimulation protocols that are typically utilized while collecting SEMR data in the intact and injured spinal cord in rats and humans (Minassian *et al*., 2004; Gerasimenko *et al*., 2006; Lavrov *et al*., 2006; Capogrosso *et al*., 2013; Gad *et al*., 2015; Hofstoetter *et al*., 2018). The stimulation procedure and protocol was similar to as described previously (Sharma & Shah, 2021b). Briefly, rats were placed in an acrylic cylinder and allowed to calm down for 2-3 minutes before connecting the head connector to the amplifier’s differential cable. Two bipolar (C6(+)C8(-) and C6(-)C8(+)) epidural electrode configurations were used to deliver ES. Data was collected at multiple time points across three months with each session lasting between 20 minutes to 5 hours. The total duration of stimulation was approximately 17 hours and included the following stimulation protocols (**Figure 1-b**):

1. **Single-pulse cervical ES:** A single-pulse stimulation protocol is usually employed to characterize the obtained evoked responses and involves ES at different stimulus intensities (Gerasimenko *et al*., 2006; Sharma & Shah, 2021b). In our experiment, the stimulation intensity ranged from 100-800 μA, in steps of 100 μA, and a single stimulation pulse was provided every three seconds. A total of 30-40 stimulation pulses were delivered for each stimulation intensity, and a rest period of no stimulation was provided for 60 seconds between consecutive stimulation intensities.
2. **Paired-pulse cervical ES:** Paired-pulse stimulation protocol are typically employed to investigate the physiological status of intact and injured spinal cord by measuring the extent of post-synaptic activation depression (Hultborn *et al*., 1996; Gerasimenko *et al*., 2006). Paired-pulse stimulation involved delivering a pair of ES with two pulses separated by a fixed inter-stimuli interval (ISI). In our experiments, the stimulation intensity ranged from 100 μA to 800 μA in steps of 100 μA. The frequency of paired-pulse stimulation was 0.3 Hz, and the inter-stimuli interval was 10, 30, 50, 100, 500, 1000, and 2000 msec. A total of 12-15 stimulation pulses were delivered for each stimulation intensity and ISI. A 60-second rest period was allowed between consecutive ISIs at the same stimulation intensity.
3. **Multiple frequency cervical ES:** Similar to paired-pulse ES, multiple frequency stimulation protocols are typically employed to investigate the physiological status of intact and injured spinal cord by measuring the extent of post-synaptic activation depression (Lloyd & Wilson, 1957; Lavrov *et al*., 2006). In our experiment, stimulation was delivered at frequencies of 0.2, 0.5, 1.0, 5.0, 10.0 and 30.0 Hz at the minimum stimulation intensity that evoked robust responses in all the implanted muscles. A total of 12-15 stimulation pulses were delivered at each stimulation frequency. Between every stimulation frequency, a rest period of no stimulation was provided for sixty seconds.

All cervical SEMR EMG signals were filtered (band passed, 10 Hz-5 kHz) and amplified (x1000) using an analog amplifier (differential AC amplifier, AM-systems Inc., USA). The 60Hz filter notch was on. The signal was digitized at a 10 kHz sampling rate using a data acquisition unit (LabChart, AD Instruments). ES was performed using a Grass S88 Stimulator (Grass Instruments) that outputted square-wave monophasic with a fixed pulse width of 200 µsec. To obtain constant-current stimulation, a stimulus isolation unit (Grass SIU-C, Grass Instruments) in series with the Grass stimulator was used. The stimulation current intensity was monitored continuously by measuring the voltage drop across a 1KOhm resistor in series with the rat using the digital input of the data acquisition unit. All SEMR data collection sessions accompanied simultaneous video recording of the rat’s functional behavior with a LabChart integrated camera or 60 frames per second (fps) video camera. This was important to discard later any EMG data that might be contaminated with irrelevant forelimb activity. Details of experimental procedures are provided below.

### Cervical SEMR: baseline and 100d

Cervical SEMR secondary to single-pulse bipolar ES were collected before (baseline) and end of the whole stimulation period (100d). For baseline, different stimulus intensities (100-800 μA, in steps of 100 μA) with appropriate rest periods between intensities were delivered using bipolar configurations. During 100d recordings, except for one rat, SEMR was collected only at 400 µA (minimum stimulation required to evoke robust SEMR) and 800 µA (maximum allowable stimulation intensity). A total of 30-40 responses per intensity per configuration were collected during both recording sessions.

### Reaching & grasping: baseline and 100d

Reaching & grasping was collected before (baseline) and end of the whole stimulation period (100d). All rats were re-trained to achieve a seventy percent success rate on reaching & grasping before the final recording. For both recording sessions, rats were placed individually in a reaching & grasping chamber, and a total of ten successful reaching & grasping events along with EMG from the implanted muscles were collected. The whole testing procedure was video recorded using a 60 frames per second (fps) camera. A battery-powered pulse generator was used to produce a TTL pulse that helped synchronize the reaching & grasping video and EMG data (Rascoe *et al*., 2018).

### Data analysis

Only SEMR data secondary to single-pulse stimulation before (baseline) and end of the whole stimulation period (100d) was included in the data analysis. Repeated stimulation during SEMR data collection between baseline and 100 day acted as an experimental variable and the obtained SEMR were not analyzed. Baseline and 100 day cervical SEMR data were analyzed using a custom-built MatLab program. The responses obtained during the procedures were extracted and assessed for signal quality. Responses that appeared noisy or aberrant, corresponded to irrelevant physical activity in the testing apparatus, and resulted from the stimulation pulse(s) higher or lower than the selected stimulation intensity were rejected and excluded from the final analysis. For analysis, a 35 msec window was selected for each SEMR: 5 msec before the onset of stimulation was considered as the baseline for that signal, and 30 msec after the stimulation was the actual evoked response to the stimulation. Evoked responses within this 30 msec window of stimulation were divided into three types: early (ER), middle (MR) and late (LR) as mentioned here (Sharma & Shah, 2021b). For further analysis, SEMR were rectified by taking absolute of their values. Mean of the obtained responses at the same stimulation intensity was used to examine the raw peak-rectified amplitudes for ERs and MRs, and integrated EMG (iEMG) for LRs. For ER and MR, the amplitude difference between the baseline and the peak of the response was defined as the peak-rectified amplitude. For LRs, the obtained iEMG (area under the curve using trapezoidal numerical integration) was divided by the duration of the responses (10 msec to 30 msec = 20 msec) to obtain average iEMG. To identify the stimulation intensities at which the raw amplitudes of the obtained responses demonstrate least variations, we calculate the segmental variance. For segmental variance, points of distinct change in the series of a raw amplitude data were determined first by iteratively minimizing the sum of the cost functions of each segment between potential change points. Later, the segmental variance is calculated as the variance between the two consecutive points of distinct change (MathWorks, 2020).

To obtain normalized peak-rectified amplitude and iEMG, the peak-rectified amplitudes of ER and MR, and average iEMG at 400 µA were normalized to the peak-rectified amplitude of ER at 800 µA. We chose peak-rectified amplitude of ER as the raw amplitude of these response plateaus and demonstrated stable segmental variances at higher stimulation intensity (Sharma & Shah, 2021b).

For reaching & grasping, only the trials that started with the rat facing the anterior wall of the cage and the paw on the floor, retrieved the pellet in a single take, and had associated noise or artifact-free EMG signals were included in the final analysis (8-10 trials/rat). To account for the difference in the time duration of multiple reaching & grasping trials within and between rats, duration of the reaching & grasping trials was represented between 0-100 % using cubic spline interpolation, where 0 and 100% indicated the start and end of a trial, respectively. EMG activity that corresponded to the reaching & grasping trials were band-passed filtered (20-1000 Hz) using 4^th^ order Butterworth filter, in forward and reverse direction. The filtered data was rectified by taking the absolute of their values and area under the curve (integrated EMG, iEMG) from the R&G trials. To calculate normalized iEMG, the obtained iEMG was normalized to the duration of the R&G event.

### Statistical analysis

Wilcoxon test was used to perform statistical analysis for the difference in the raw amplitudes and iEMG of the obtained SEMR, and normalized iEMG during R&G for the baseline and at 100d recordings. All statistical analyses were performed using SPSS (IBM Corporation, Armonk, NY, USA) and MatLab (MathWorks Inc., Natick, MA, USA). Statistical significance threshold was set at 0.05.

## Results

### Cervical SEMR: baseline vs. 100d

We characterized cervical spinal evoked motor responses (SEMR) secondary to epidural stimulation (ES) as early (ER), middle (MR), and late responses (LR) as described recently (Sharma & Shah, 2021b). ER appeared at a relatively higher stimulation intensity with latencies between 1-3 msec. In contrast, MR and LR appeared at a relatively lower stimulation intensity with latencies between 3-5 and 9-11 msec, respectively (**Figure 2)**. In contrast to the baseline cervical SEMR, all three responses were elevated at the end of 100d after ∼17 hours of stimulation sessions, in all muscles (only two muscles shown in **Figure 2**).

**Figure 2:**
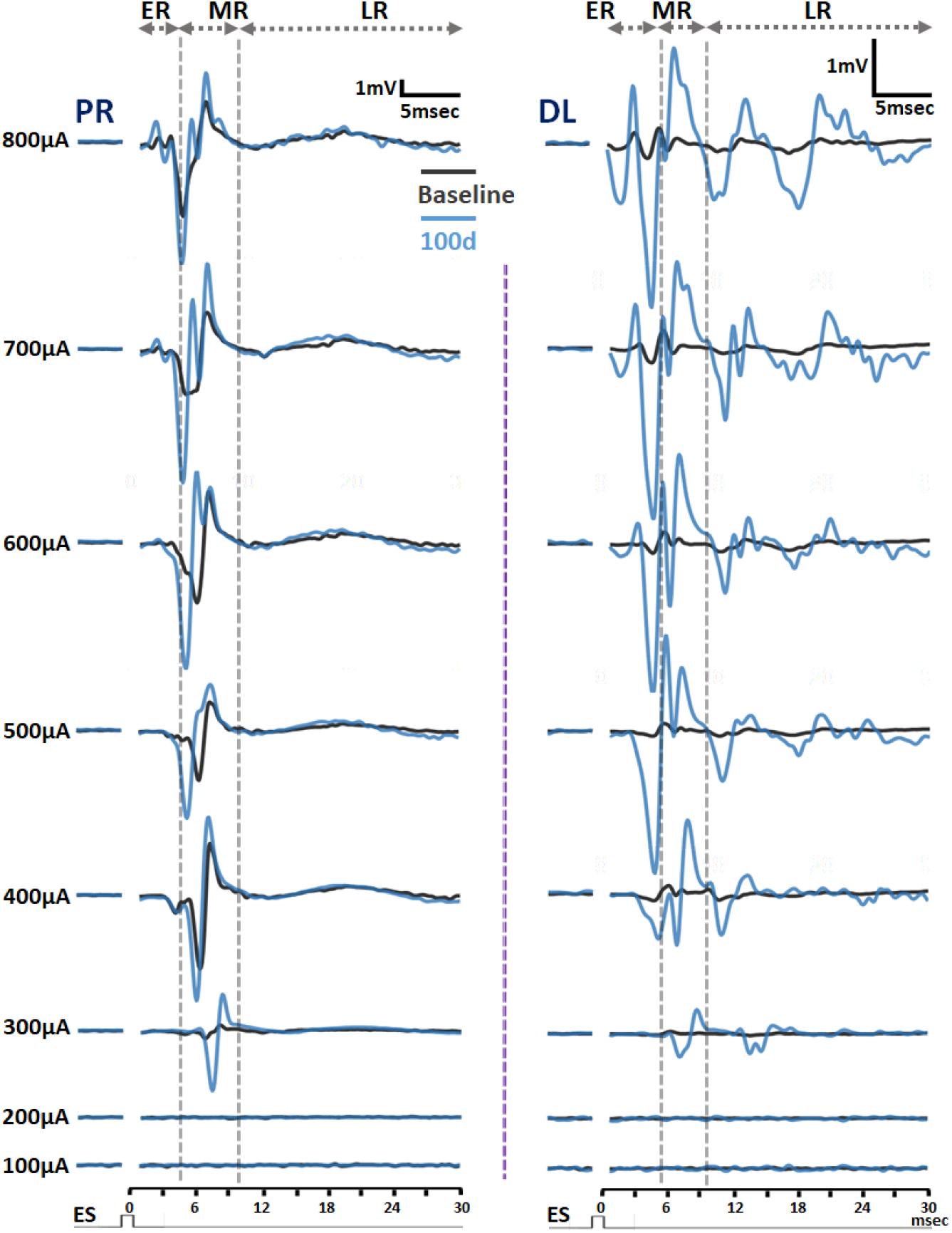
Cervical spinal evoked motor responses (SEMR) baseline vs. 100d: SEMR evoked in two forelimb muscles (PR and DL) secondary to single-pulse ES at baseline (black traces) and at 100d (blue traces) at different stimulation intensities in a representative rat. Each trace is an average of 25-35 individual responses. ER, MR, and LR regions are represented within vertical grey dashed lines. Note increased amplitude of the obtained responses during final recording as compared to the initial recording. PR: pronator; DL: deltoid; ER: early responses; MR: middle response: LR: late response.

The facilitation was more prominent at higher stimulation intensities than the lower. Additionally, the stimulus-threshold, defined as the minimum current required to elicit an evoked response, were relatively lower at 100d compared to at the baseline (300-400 µA on 100d vs. 400-600 µA at baseline for DL in **Figure 2**). Once elicited at the stimulus-threshold intensity, greater increases in the magnitude of the evoked responses with increased stimulation intensity was observed for 100d, indicating greater recruitment of sensory afferents and motor axons that eventually engages more muscle fibers to contract (see discussion). Additionally, muscles that failed to evoke a response at baseline testing elicited a response at 100d (in **Figure 3**, note the absence of LR in FDS at baseline).

**Figure 3:**
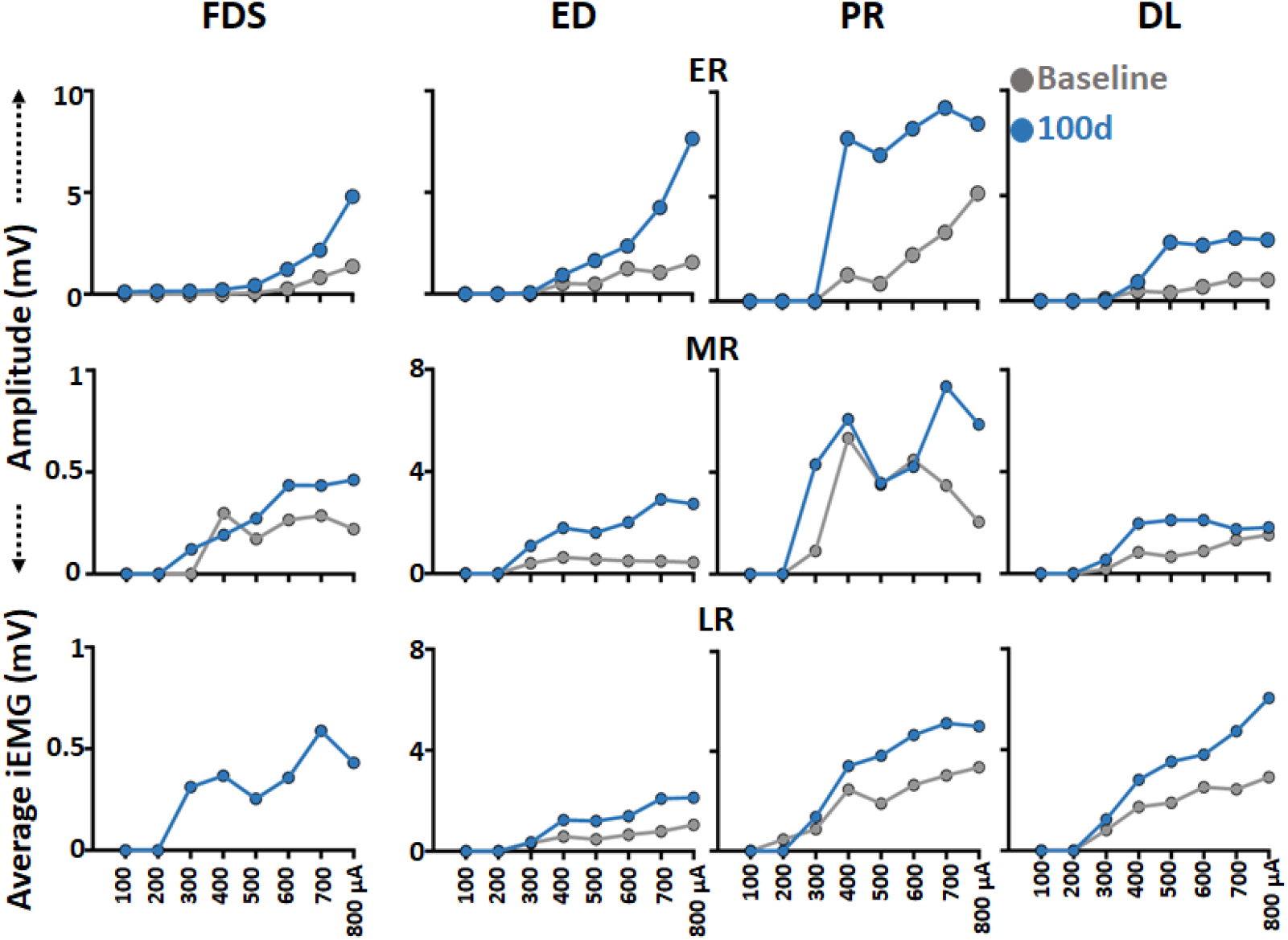
Stimulus-response baseline vs. 100d: Stimulus-response curves of ER, MR. and LR from the four forelimb muscles (FDS, ED, PR, and DL) during single-pulse bipolar ES in a representative rat at baseline and 100d. Note, persistently increased amplitude of ER and MR, and iEMG of LR on 100d compared to at the baseline. FDS: flexor digitorum superficialis; ED: extensor digitorum; PR: pronator; DL: deltoid; ER: early responses; MR: middle response: LR: late response.

When plotted for all muscles together at 400 and 800 µA, the obtained response amplitudes at 100d were significantly higher than the baseline recording. The increase was more robust for MRs and LRs as compared to ERs. Specifically, at 100d, the peak-rectified MR amplitudes for FDS (z = -2.201, p = 0.028), ED (z = -2.201, p = 0.028), PR (z = -2.201, p = 0.028), DL (z = -2.201, p = 0.028) at 400 µA, and ED (z = -1.992, p = 0.046), PR (z = -2.201, p = 0.028), DL (z = -2.201, p = 0.028) at 800 µA were significantly higher than at the baseline. Additionally, for LRs, the average iEMG for ED (z = -2.201, p = 0.028), PR (z = -1.992, p = 0.046), DL (z = -2.201, p = 0.028) at 400 µA, and ED (z = -2.201, p = 0.028), DL (z = -2.201, p = 0.028) at 800 µA were significantly higher at 100d compared to at the baseline recording (**Figure 4**). The raw peak-rectified ER amplitudes were significantly higher only for FDS at 400 µA (z = -1.992, p = 0.046) and DL at 800 µA (z = -2.201, p = 0.028). We did not observe significant increase in the peak-rectified MR amplitudes for FDS (z = -1.572, p = 0.116) at 800 µA, average iEMG for FDS at 400 µA (z = -1.826, p = 0.068) and FDS (z = -1.826, p = 0.068) and PR (z = -1.572, p = 0.116) at 800 µA at 100d compared to at the baseline recording.

**Figure 4:**
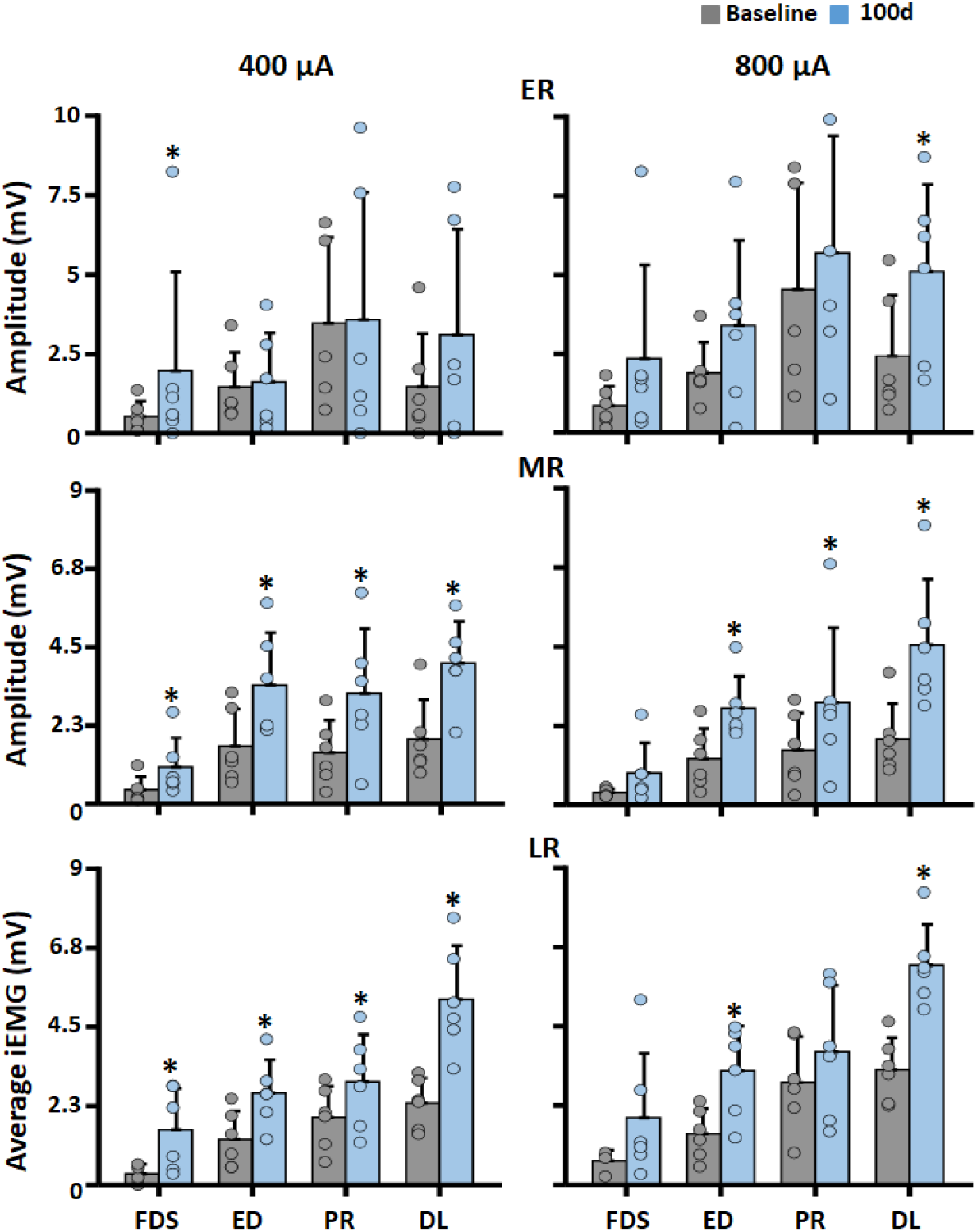
Raw peak-rectified amplitude and iEMG baseline vs. 100d: Bar graphs for raw peak-rectified amplitudes of ER and MR, and iEMG for LR during single-pulse bipolar ES at 400 and 800 µA from four different muscles at the baseline and 100d. Color dots within each bar graph represents data from individual rats. *p<0.05, significantly different baseline. FDS: flexor digitorum superficialis; ED: extensor digitorum; PR: pronator; DL: deltoid; ER: early responses; MR: middle response: LR: late response.

### Reaching & grasping: baseline vs. 100d

Stimulation sessions per se did not alter the reaching & grasping success rate at 100d compared to at the baseline. However, in comparison to the baseline, all muscles demonstrated an increase in raw amplitudes at 100d (**Figure 5-a**).

**Figure 5:**
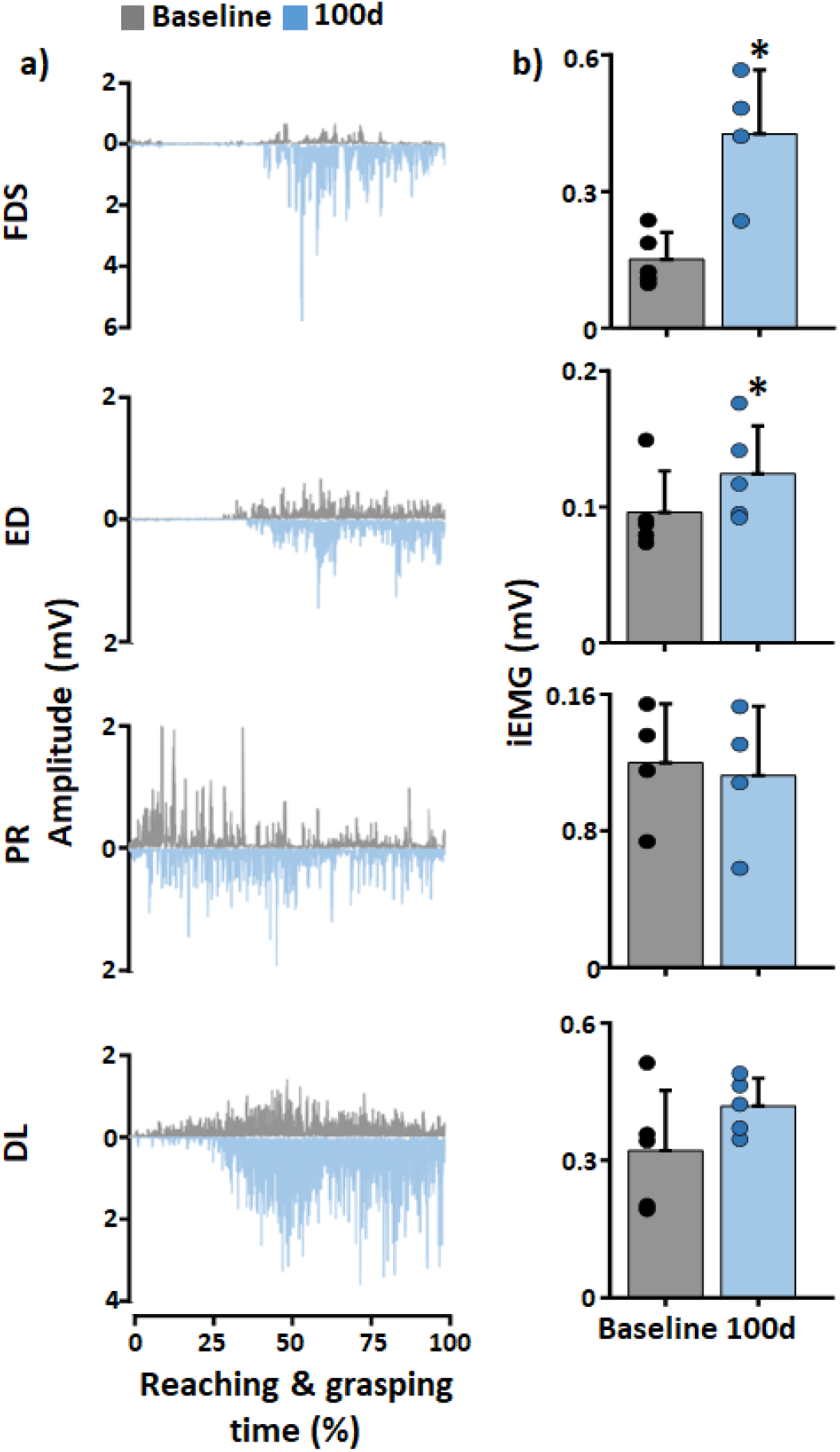
Reaching & grasping baseline vs. 100d: **a)** Shown are the rectified EMG of various forelimb muscles (FDS, ED, PR, DL) during Reaching & grasping task in a representative rat at the baseline and 100d. **b)** Bar graphs showing normalized iEMG during reaching and grasping during baseline vs. 100d. Color dots within each bar graph represents data from individual rats. *p<0.05, significantly different baseline. FDS: flexor digitorum superficialis; ED: extensor digitorum; PR: pronator; DL: deltoid; iEMG: integrated EMG.

When plotted for all rats together there was a statistically significant increase in iEMG for the FDS and ED muscles at 100d compared to the baseline (FDS: z = -2.023, p = 0.043; ED: z = -2.023, p = 0.043; PR: z = -18.26, p = 0.068; DL: z = -1.483, p = 0.138) (**Figure 5-b**), providing evidence for an increased muscle power production during reaching & grasping post repeated stimulation during SEMR data acquisition. Additionally, significant increase in muscle activity was only seen for the distal muscles, FDS and ED, with motoneuron pool spanning across C6-C8 compared to PR and DL with motoneuronal pool mainly located around C4-C6 signal segment (McKenna *et al*., 2000; Tosolini & Morris, 2012). The finding indicating that the effects of repeated ES are more prominent for the muscles whose motoneurons are more exposed to the ES induced electric field.

### Cervical SEMR: normalized baseline vs. 100d

As the raw amplitudes of the obtained SEMR may vary within and between rats, especially when the evoked responses are collected longitudinally, normalizing these responses to a stable component is often recommended to allow meaningful comparisons of the obtained responses. To identify the stable component(s), we investigated the change in raw peak-rectified amplitudes and associated segmental variance at different stimulation intensities **(Supplementary Figure 1-a)**. As seen in the figure for ED muscle in three representative rats, at 600 µA and higher stimulation intensity, we did not see any abrupt change in the segmental variance of ER and LR, indicating lower variability in the amplitude of the responses at higher stimulation intensities. In contrast, raw amplitude and segmental variance of MRs at same stimulation intensities either increased, decreased or remained relatively unchanged. Segmental variance of ERs, MRs, and LRs obtained at different stimulation intensities for ED for all rats indicated that the segmental variance of all the components increased until 100 to 500 µA **(Supplementary Figure 1-b)**. At higher stimulation intensities (600-800 µA), the segmental variance for ER and LR remained relatively stable. In contrast, for MR, there were trends of reduced segmental variance at 700 and 800 µA compared to the 500 µA. Similar trends for segmental variance were noted for different components of cervical SEMR at higher stimulation intensities for other muscles. Although segmental variance for ER and LR behaved in a similar manner at higher stimulation intensities, we decided to normalize the obtained responses with peak-rectified amplitude of ER as these responses are secondary to the activation of motor axons (no modifiable synaptic involvement) and less complex than LRs that are known to be secondary to the activation of sensory afferent innervating a complex set of interneurons (Gerasimenko *et al*., 2006; Capogrosso *et al*., 2013; Sharma & Shah, 2021b). Additionally, it was easier to identify single peak-rectified amplitude for ER as they were phase-locked, synchronous in nature, and had relatively stable amplitudes compared to other responses. In contrast, LRs consisted of multiple peaks that were asynchronous in nature (not time and phase-locked) (Sharma & Shah, 2021b). On normalizing the obtained raw amplitudes or iEMG at 400 µA with peak-rectified amplitudes of ER at 800 µA, we did not see significant difference between the responses at 100d compared to at the baseline for ER (FDS: z = -1.153, p = 0.249; ED: z = -0.314, p = 0.753; PR: z = -0.405, p = 0.686; DL: z = -0.314, p = 0.753), MR (FDS: z = -0.734, p = 0.463; ED: z = -1.992, p = 0.046; PR: z = -0.674, p = 0.500; DL: z = -0.524, p = 0.600), and LR (FDS: z = -0.365, p = 0.715; ED: z = -1.572, p = 0.116; PR: z = -0.674, p = 0.500; DL: z = -0.105, p = 0.917) at 400 µA and MR (FDS: z = -0.105, p = 0.917; ED: z = -1.572, p = 0.116; PR: z = -0.135, p = 0.893; DL: z = -0.105, p = 0.917), and LR (FDS: z = -0.734, p = 0.463; ED: z = -1.153, p = 0.249; PR: z = -0.405, p = 0.686; DL: z = -0.734, p = 0.463) at 800 µA (**Supplementary Figure 2**).

## Discussion

Data from this work demonstrate that cervical spinal evoked motor responses (SEMR) at rest and electromyographic (EMG) activity during reaching & grasping task are modulated in adult non-injured rats secondary to repeated epidural stimulation (ES) during SEMR data acquisition spanning across 100 days for ∼17 hours. Below we discuss the observed changes, potential mechanisms resulting in these changes, and precautions to be practiced while drawing functional interpreting from the SEMR data.

### Increased spinal cord and cortical excitability

The observed facilitation in cervical SEMR was more robust for the monosynaptic middle (MR) (physiologically similar to H-reflex) and polysynaptic late responses (LR) as compared to the direct motor early responses (ER) (physiologically similar to M-wave). Several *in-vitro* and *in-vivo* studies involving spinal cord and peripheral nerve electrical stimulation have demonstrated similar adaptations following short-duration stimulation. In healthy humans, transcutaneous cervical and lumbar spinal cord stimulation for 20-30 minutes is known to facilitate spinal and cortical evoked responses with effects lasting for an hour post-stimulation (Benavides *et al*., 2020; Kumru *et al*., 2021). Similar electrophysiological adaptations are also seen in adult non-injured rats secondary to lumbar ES for 15 minutes resulting in increased ER and MR amplitude with effects lasting for approximately 8 minutes post-stimulation (Taccola *et al*., 2020). Similarly, patterned stimulation of tibial and median nerve results in increased monosynaptic response activity lasting for approximately for 1-16 minutes post-stimulation (H-reflex) without causing significant changes in the direct motor responses (M-wave) (Kitago *et al*., 2004; Yeh *et al*., 2015; Pearcey & Zehr, 2020). *In-vitro* tetanic stimulation of the neonatal spinal cord dorsal horn for 30 min-2.5 hours in rats results in increased evoked potential amplitudes recorded from the ventral horn lasting for ∼2.5 hours post-stimulation (Pockett & Figurov, 1993). Moreover, facilitation of monosynaptic response recorded from the ventral roots in cats is reported following 12 seconds of gastrocnemius nerve stimulation (Lloyd, 1949).

In addition to SEMR, we also observed increased EMG output from forelimb muscles during the reaching & grasping task without any changes in quality of performance. An increase in the EMG output secondary to cervical transcutaneous stimulation along with exercise training is commonly reported in healthy humans (Kumru *et al*., 2021). However, in the present experiments similar neuromodulatory changes were observed in non-injured adult rats receiving ES alone, indicating spinal and cortical plastic changes secondary to short-duration stimulation in the absence of motor training (Yamaguchi *et al*., 2020).

### Potential mechanisms contributing to increased excitability

The prominent facilitation of MRs and LRs can be attributed to increased excitability of the predominant structures activated with cervical ES at lower stimulation intensities, namely the sensory afferent fibers (Gerasimenko *et al*., 2006; Capogrosso *et al*., 2013). Indeed, Jiang *et al*. (2016) demonstrated that electrical stimulation of the sensory afferent supplying forelimb muscles for 6hours/day for 10 days significantly increases the sensory afferent density in the dorsal column and afferents fiber arborization onto spinal motoneurons and interneurons in non-injured adult rats. Although not as intensive in stimulation duration, stimulation for only 17 hours spread across five sessions and three months was enough to induce plasticity in our experiments.

Repeated stimulation of the sensory afferent increases the size of endplate synaptic potentials (EPSP) and activation of synaptic buttons, which in turn increases neurotransmitter release from the synaptic vessels (Eccles & Krnjevic, 1959; Lev-Tov *et al*., 1983). Greater availability of the neurotransmitter results in increased activation of motoneurons and might play a crucial role in the observed facilitation of SEMR and EMG activity secondary to repeated stimulation (Purves D, 2001). EPSP potentiation also increases neurotransmitter release at the neuromuscular junction and effectively increases cross-linkages during muscle contraction and, hence, increased EMG output (Kandel *et al*., 2012). Additionally, repeated stimulation also decreases the pre-synaptic inhibition (Kaczmarek *et al*., 2017; Pearcey *et al*., 2017) and facilitates the formation of new connections of sensory afferents with motoneurons and interneurons (Jiang *et al*., 2016). These additional connections might be responsible for the observed facilitation of MRs and LRs secondary to repeated stimulation in our experiments.

In contrast to MRs and LRs, ERs originate secondary to the activation of motor axons at higher stimulation strengths (Lavrov *et al*., 2006; Sharma & Shah, 2021b). We observed trends of increased amplitudes at the end of the stimulation period, indicating towards increased excitability of the motor axons. Most likely, the duration of stimulation was not enough to establish significant plastic changes in the motor axons. Compared to sensory afferents, motor axons exhibit a lesser number of slow inactivating Na+ channels in humans and rats, implying a lesser influx of Na+ into motor axons before their deactivation (Honmou *et al*., 1994; Bostock & Rothwell, 1997). Lesser intracellular Na^+^ results in low amplitude motor evoked responses (reviewed in Bostock & Rothwell, 1997). Indeed, M-wave originating from motor axons, similar to ER, fail to demonstrate significant facilitation indicating the absence of neuro-myal potentiation (Kitago *et al*., 2004; Yeh *et al*., 2015). In addition to the spinal cord, mechanisms acting at the cortical level, such as increased excitability of the corticospinal projections and reduced GABAergic activity resulting in long-lasting facilitatory changes, cannot be ignored (Kaelin-Lang *et al*., 2002; Golaszewski *et al*., 2012).

In our experiment, the discussed adaptations are most likely cumulative in nature, with each stimulation session likely causing the above-mentioned short-term physiological changes, and with repeated stimulation, permanent changes occur.

### Precautions while interpreting SEMR data

Raw amplitude provides an appropriate estimate of the ongoing physiology, especially in the studies involving chronic intramuscular implants that is less affected by the variables such as electrode position during repeated testing (Palmieri *et al*., 2004). However, reporting raw amplitudes fails to account for intrinsic variables, such as differences in the muscle size and motor unit approximation to recording electrode, restricting pooling data from multiple participants and warrants involving normalization procedure. Based on the reported data, we recommend exercising precautions while normalizing SEMR component with the least variable ER (Lavrov *et al*., 2006) component as normalization may mask the real physiological adaptations secondary to the repeated stimulation, as shown in supplementary figure 2.

Although elevated EMG amplitudes suggest increased spinal cord and/or cortical excitability following ES (Angeli *et al*., 2014; Rejc *et al*., 2015; Capogrosso *et al*., 2016; Asboth *et al*., 2018), SEMR may not necessarily accompany the observed motor changes. For example, increased MR amplitudes following thoracic transaction has been demonstrated in spontaneously recovering rats that failed to recover bipedal stepping and in rats demonstrating improved bipedal treadmill stepping secondary to motor training and lumbar ES (Lavrov *et al*., 2006; Courtine *et al*., 2009). Indeed, elevated spinal cord excitability evident from the SEMR amplitudes may not be enough to result in improved functional outcomes (Lavrov *et al*., 2006). Therefore, correlating the obtained evoked responses with appropriate motor behavior measures for functional interpretation is a common practice and SEMR data are rarely used alone. In rats, restoration of the treadmill walking following thoracic transactions and improvement in reaching & grasping following a C5 dorsal crush correlates with the re-emergence of LRs (Lavrov *et al*., 2006; Lavrov *et al*., 2008) and gradual change of the SEMR profile towards pre-injury status (Alam *et al*., 2017), respectively. Similarly, increased ERs and decreased LRs in humans receiving orthosis-assisted ambulatory training on the treadmill following a SCI correlates with decreased muscular exhaustion and better performance during assisted stepping (Dietz *et al*., 2009; Hubli *et al*., 2012).

### Therapeutic implications

As repeated ES during SEMR data acquisition can itself induce physiological adaptations in the spinal cord and/or brain evident from the increased SEMR and forelimb EMG activity, we recommend accounting for the SEMR data acquisition duration and employing stimulation protocols essential to investigate underlying physiology. Additionally, sole reliance on SEMR profiles for functional recovery interpretation may not be a pragmatic approach and including a motor behavior correlate is warranted. Finally, since most SCIs are incomplete in nature, plastic changes seen in the intact spinal cord following repeated ES can be expected to happen in an injured spinal cord too. Hence, increased spinal cord excitability and improved muscle force production can be potential recovery mechanisms with repeated ES following a SCI (Angeli *et al*., 2014; Rejc *et al*., 2015; Lu *et al*., 2016; Alam *et al*., 2017).

## Limitations

Including control rats not receiving ES would have allowed investigating the natural changes in SEMR and EMG profiles resulting from the rat’s aging. However, our rats were 20-33 weeks old, normally regarded as adults, and we do not expect any changes in physiology (Sengupta, 2013).

## Conclusion

Overall, our study demonstrates that repeated ES during SEMR data acquisition alone can facilitate the SEMR at rest and EMG during forelimb motor tasks in non-injured adult rats. The findings indicate potential plastic changes at the spinal and cortical level secondary to repeated ES, and advocate practicing cautions, such as accounting for the duration of SEMR data acquisition, including appropriate control groups and normalization methods, and motor behavior correlate, while utilizing SEMR as a biomarker of recovery in longitudinal studies involving repeated SEMR data acquisition.

## Acknowledgments

This work was supported by the Craig Neilsen Grant # 338237 granted to P.K.S and New York State Pre-Doctoral Fellowship Grant # DOH01-C33614GG-3450000 granted to P.S.

## Supplementary Data

**Supplementary Figure 1:**
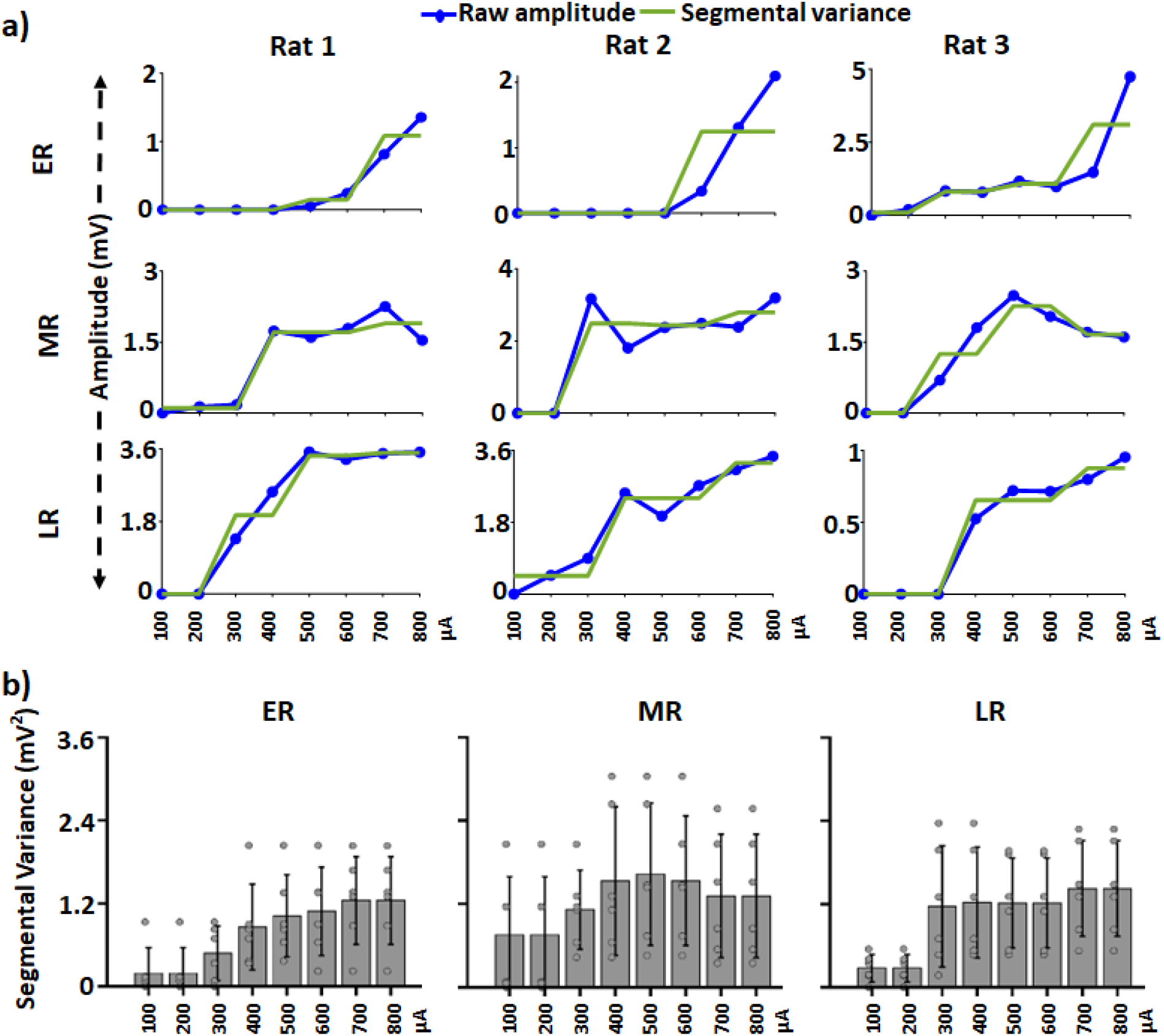
Raw amplitude and segmental variance for cervical spinal evoked motor responses (SEMR) at baseline: **a)** Raw amplitudes (blue) and segmental variance (green) of ER, MR, and LR for ED during single-pulse evoked responses in three representative rats at baseline. **b)** Bar graphs for the segmental variance of ER, MR, and LR for ED during single-pulse bipolar ES at baseline. Gray dots within each bar graph represents data from individual rats. ED: extensor digitorum. ER: early responses; MR: middle response: LR: late response.

**Supplementary Figure 2:**
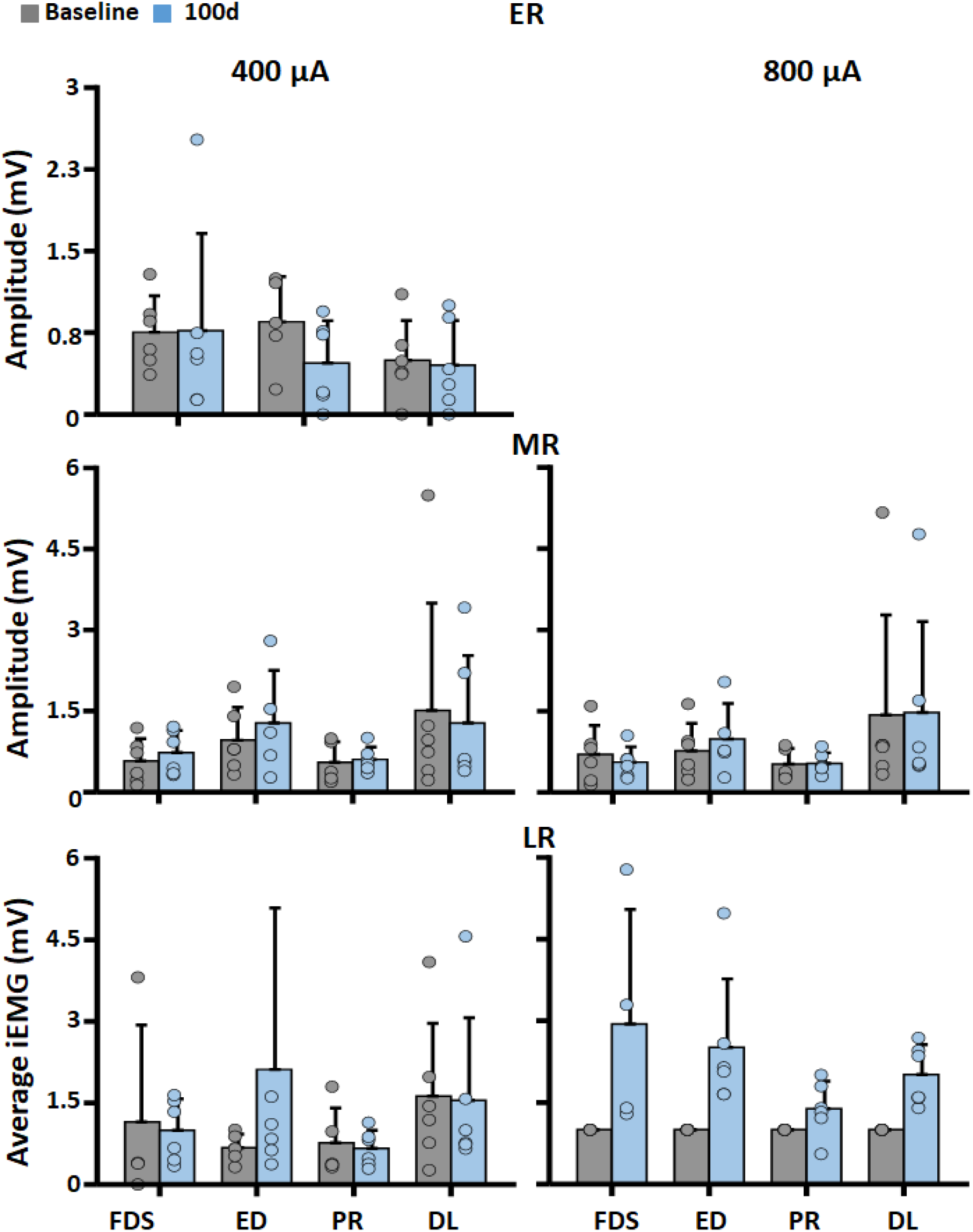
Normalized peak-rectified amplitude and iEMG baseline vs. 100d: Bar graphs for normalized peak-rectified amplitudes of ER and MR, and iEMG for LR during single-pulse bipolar ES at 400 and 800 µA from four different muscles at the baseline and 100d. Color dots within each bar graph represents data from individual rats. FDS: flexor digitorum superficialis; ED: extensor digitorum; PR: pronator; DL: deltoid; ER: early responses; MR: middle response: LR: late response. Note: No bar graph for ER at 800 µA as all amplitudes will be equal to 1 mV (ER at 800 µA/ ER at 800 µA).

